# HIV Infection and Opioid Treatment Enable the Engraftment of Kaposi Sarcoma-like Tumors into Immunocompetent Mice

**DOI:** 10.1101/2025.06.30.662265

**Authors:** Julian Naipauer, Ezequiel Lacunza, Anuj Ahuja, Santas Rosario, Carolina Alejandra Alvarez Canete, Martin C. Abba, Ethel Cesarman, Sundaram Ramakrishnan, Enrique Mesri, Sabita Roy, Umakant Sharma

## Abstract

The Kaposi Sarcoma herpesvirus (KSHV) causes Kaposi sarcoma (KS), primary effusion lymphoma, a lymphoproliferative disease (KSHV-multicentric Castleman’s disease), and a cytokine inflammatory syndrome (KICS). These diseases occur more frequently, though not exclusively, among people living with HIV or other types of immune dysregulation. While limited KS can regress with immune reconstitution, such as through antiretroviral therapy (ART) in people living with HIV, there are currently no curative treatments for advanced KS. Preventive or therapeutic vaccines targeting KSHV could have a significant clinical impact; however, the development and testing of such strategies have been limited by the lack of preclinical models that faithfully recapitulate KS, including the presence of infected spindle cells and a relevant immune microenvironment. HIV/AIDS is an important cofactor for KS, and globally, the majority of individuals with KS are HIV-infected. Current evidence indicates that HIV-1 may enable KS progression through immunosuppression and promote pathogenesis by inducing inflammatory cytokines and producing secreted regulatory proteins like Tat and Nef. The design and testing of new therapeutic approaches based on pathogenesis are hampered by the lack of models that replicate KSHV oncogenesis in the context of HIV/AIDS. In the present study, we demonstrate that KSHV-infected cells can form tumors in an immunocompetent mouse model, but only when these mice are pretreated with EcoHIV and morphine, both of which have immunomodulatory and pro-inflammatory effects. These tumors exhibit gene expression profiles and immune microenvironments that closely resemble those observed in human KS lesions. This novel KSHV tumor model in immunocompetent mice provides a valuable platform to test immunotherapeutic strategies for KS, including immunomodulatory agents, targeted antibody therapies, checkpoint inhibitors, and vaccines.

**Author Summary:** Kaposi sarcoma (KS) is a cancer caused by the Kaposi’s sarcoma-associated herpesvirus (KSHV), also known as human herpesvirus 8 (HHV-8). A vaccine against this virus could prevent or cure this malignancy. However, there are no good animal models with an immune system to test new immunization approaches. The possibility of having KSHV-driven tumors develop in immunocompetent mice due to EcoHIV infection presents a unique opportunity to create an AIDS-KS mouse model suitable for testing immune-based therapies and KSHV vaccines in a scenario that replicates important aspects of AIDS-KS. These first-in-kind murine models incorporating a biologically relevant mouse HIV infection tool make our models unique and original, with the potential to provide novel mechanistic and pathobiological insights.

## Introduction

Kaposi’s sarcoma (KS) is an AIDS-associated malignancy (HIV/AIDS-KS) caused by the KS herpesvirus that remains a major global health challenge[1, 2]. KS can be treated with local, anti-retroviral, and chemotherapy[2]. The incidence of AIDS-KS has decreased with the widespread use of ART among HIV-infected individuals[1, 2]. Yet, KS remains potentially life-threatening for patients with advanced or ART-resistant disease, where systemic therapy is indicated, and three FDA-approved agents that include liposomal anthracyclines are available[2-4]. Despite the effectiveness of these agents, most patients progress within six to seven months of treatment and require additional therapy[5]. While early KS can be controlled by reconstitution of an immune response with antivirals in people living with HIV (PLWH) or modulation or by medication changes in the setting of iatrogenic KS, these approaches are insufficient in advanced KS or not possible in people with KS without any apparent immunodeficiency, as in the case of classic/sporadic KS. Furthermore, as patients are living longer under ART treatment, there is an increase in a new form of KS affecting patients with controlled viremia and high CD4 counts[6]. Increasing KS in PLWH with cART has been suggested as a wake-up call for research on HHV-8 [7]. The role of HIV/AIDS in KS oncogenesis includes HIV accessory molecules with pro-angiogenic and KSHV-stimulating activity and immunosuppression, which is in part mediated by immune exhaustion that could be prevented or targeted by these agents[1, 8, 9]. Currently, there is no model in which KSHV and HIV interactions can be studied that could allow for a higher understanding of the consequences of KSHV and HIV coinfection operating in HIV/AIDS-KS, nor an immunocompetent mouse model to test new immunomodulatory strategies. HIV infection is highly prevalent in injecting drug users (IDU), with upwards of 40% in various regions globally[10]. IDU has been observed to correlate with a more severe HIV pathogenesis than non-users, including much faster development of AIDS and higher rates of neurocognitive deficiencies[11,12].

Opioids, the most common IDU agent in HIV cases, have been shown in mouse models to have drastic systemic effects, including on the immune system. Opioids are the most potent analgesics and are widely used for the treatment of severe pain, such as cancer pain. However, a great number of studies have convincingly demonstrated that opioids, in particular morphine and its derivatives, are immunosuppressive. Studies in human and mouse models have suggested that one of the mechanisms by which opioids cause immunosuppression is by inhibiting IL-2 transcription in activated T lymphocytes[13, 14]. As a consequence, there are significant interactions between HIV and opioid use in the immune compartment. Recent studies indicate that compromised intestinal barrier function and subsequent microbial translocation contribute to persistent immune activation and chronic inflammation in HIV-infected patients[15, 16]. This ongoing immune activation is linked to CD4+ T-cell depletion and disease progression. Although epidemiological results are inconclusive as to whether or not opioids increase KS risk in HIV-infected individuals[17, 18], there is a need to investigate this connection, including laboratory-based evidence, given the rampant opioid epidemic that targets many HIV/AIDS-KS at-risk populations.

Currently, no mouse models reflect some important features of KS, especially the inclusion of an inflammatory tumor microenvironment which is a key consistent feature of KS lesions. Among immunocompetent mouse models, those available typically involve transgenic expression of one or few viral oncogenes; to our knowledge, the only immunocompetent model of KS with the entire KSHV is one recently reported by the Dittmer laboratory [19]. This model involved pronuclear injection of the whole viral genome, creating a transgenic mouse with all the KSHV genes, resulting in aggressive angiosarcomas in mice. This important model allows the study of different viral functions, but would not be suitable for testing immunotherapeutic strategies because the expression of viral genes during embryogenesis results in these being recognized as self, thus in tolerance and a lack of an immune response to viral antigens. Other models rely on the implantation of KSHV-infected human or mouse endothelial cells, stem cells (mesenchymal or from bone marrow), or a patient-derived xenograft into immunodeficient mouse strains[20-23]. None of these have tested the additional effects of HIV infection, in particular, controlled HIV, where there is a relatively subtle immune dysregulation, unlike that in immunodeficient mouse strains. Therefore, current KS mouse models are unsuitable for testing immunotherapies based on checkpoint inhibitors, a promising KS therapeutic modality, or KSHV vaccines.

In the present study, we developed an immunocompetent mouse model of KS, in which mouse KS-like KSHV-positive tumor growth in immunocompetent mice depends on opioid and HIV-induced immune dysregulation and/or inflammation.

## Results

### Mouse KS-like KSHV-positive tumors growing in Morphine+EcoHIV-treated immunodeficient mice engraft in immunocompetent mice

The mECK36 model of mouse KS-like KSHV-positive tumor growth was developed by transfecting KSHVBac36 into a population of bone marrow cells enriched in endothelial lineage hematopoietic cells, designated as mEC (mECK36). The injection of mECK36 into immunocompromised nude mice resulted in KSHV-bearing tumors that exhibited host and viral KS-like transcriptomes[20]. The tumorigenesis of mECK36 was associated with the upregulation of KSHV lytic genes alongside the upregulation of angiogenesis receptors and ligands, and it was dependent on the KSHV oncogene vGPCR[20]. When these tumorigenic mECK36 cells were injected into syngeneic mice, they were unable to form tumors (Supplementary Figure 1A-B).

In the opioid drug abuse population, AIDS opportunistic illnesses (AIDS-OI) remain associated with an increased risk of mortality. As opioids and HIV can be pro-inflammatory and/or immunosuppressive, we hypothesized that they could modify KSHV anti-tumor immunity and perhaps allow mouse KS-like KSHV-positive tumors to grow in immunocompetent mice. To examine this question, we evaluated the combined effects of morphine and a murine HIV infection model, EcoHIV [24]. The EcoHIV model is a genetically modified HIV that uses murine leukemia virus gp80 for cell entry in place of gp120 from HIV-1[24]. This murine leukemia virus-pseudo-typed HIV infects CD4 T cells and macrophages and can recapitulate some aspects of HIV latency, reactivation, and AIDS, and potentiates the inflammatory effects of opioids in the brain. However, unlike HIV, EcoHIV does not cause CD4 T cell depletion[25, 26]. Immunocompetent Balb/c mice were treated with saline solution, EcoHIV-infected (1×10^6^ picograms of p24), and morphine-treated (escalating dose from 15mg/kg to 25mg/kg twice a day), separately or in combination. We injected KSHV-positive mouse mECK36 cells[20, 27] (1×10^6^ cells/mouse) in the flank of saline-treated, morphine-treated, EcoHIV-infected BALB/c mice separately or in combination. Still, they did not induce tumor formation in these treated immunocompetent mice after 12 weeks post-injection (Figures 1A and 1 B). The most straightforward explanation for the growth of KSHV-infected cells in nude mice but not in syngeneic immunocompetent mice is that tumor rejection is caused by viral antigen expression leading to immunological tumor rejection. We examined the effect of morphine+EcoHIV treatments in cells injected into immunodeficient nude mice. These mice were either treated with a combination of EcoHIV (1×10^6^ picograms of p24) and morphine (escalating dose from 15mg/kg to 25mg/kg twice a day) or with saline solution and injected in the flank with tumorigenic KSHV-positive mouse mECK36 cells (1×10^6^ cells/mouse) (Figure 2A). No changes in mouse KS-like KSHV-positive tumor growth were observed between saline-treated and morphine+EcoHIV-treated nude mice (Figure 2B).

**Figure 1.**
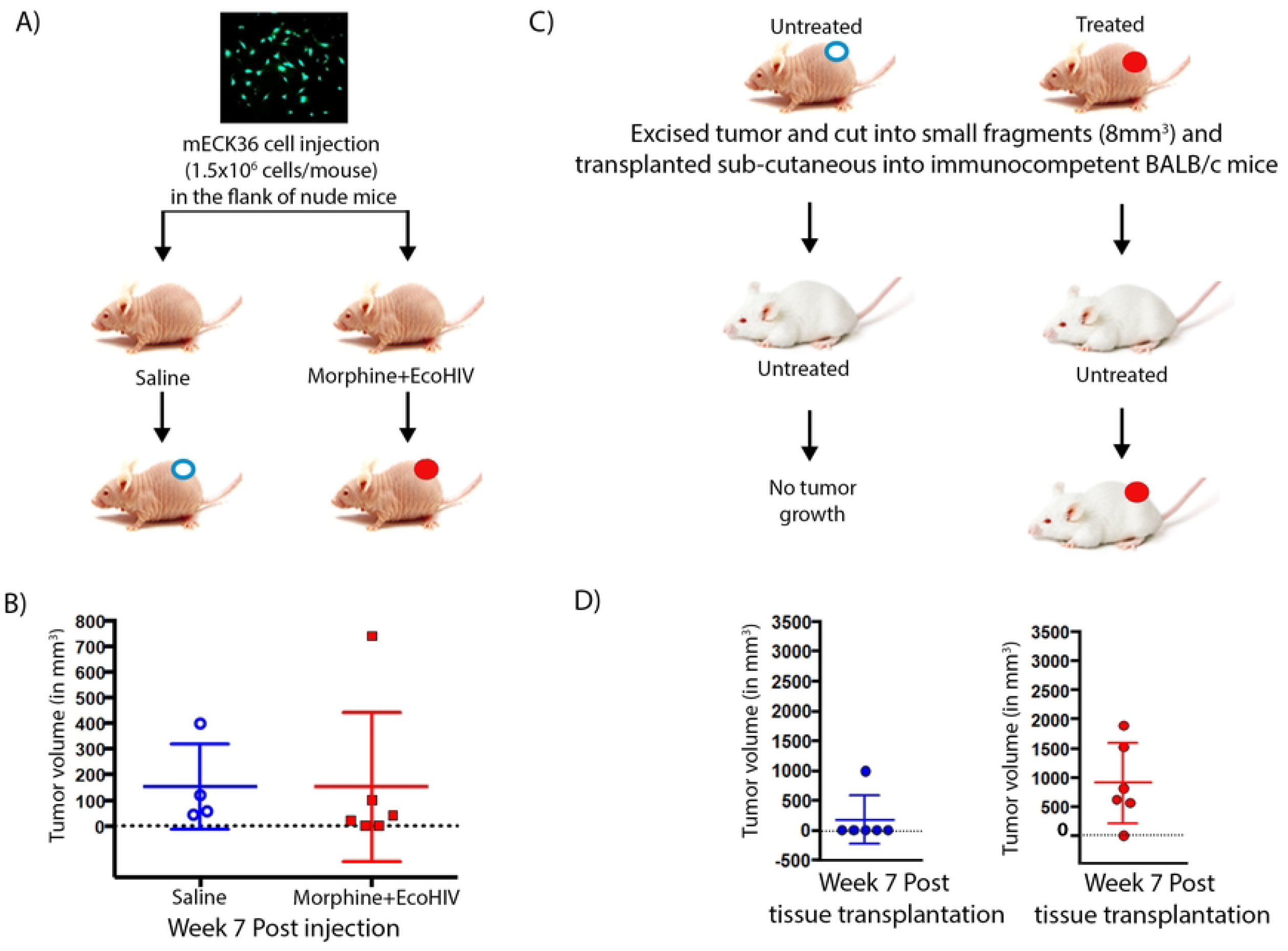
Tumorigenic mECK36 cells do not grow in immunocompetent mice treated with Morphine, EcoHIV, or a combination. A) Schematic representation of the animal injections. B) Tumor growth in Balb/c mice control, treated with Morphine, infected with EcoHIV, or the combination.

**Figure 2.**
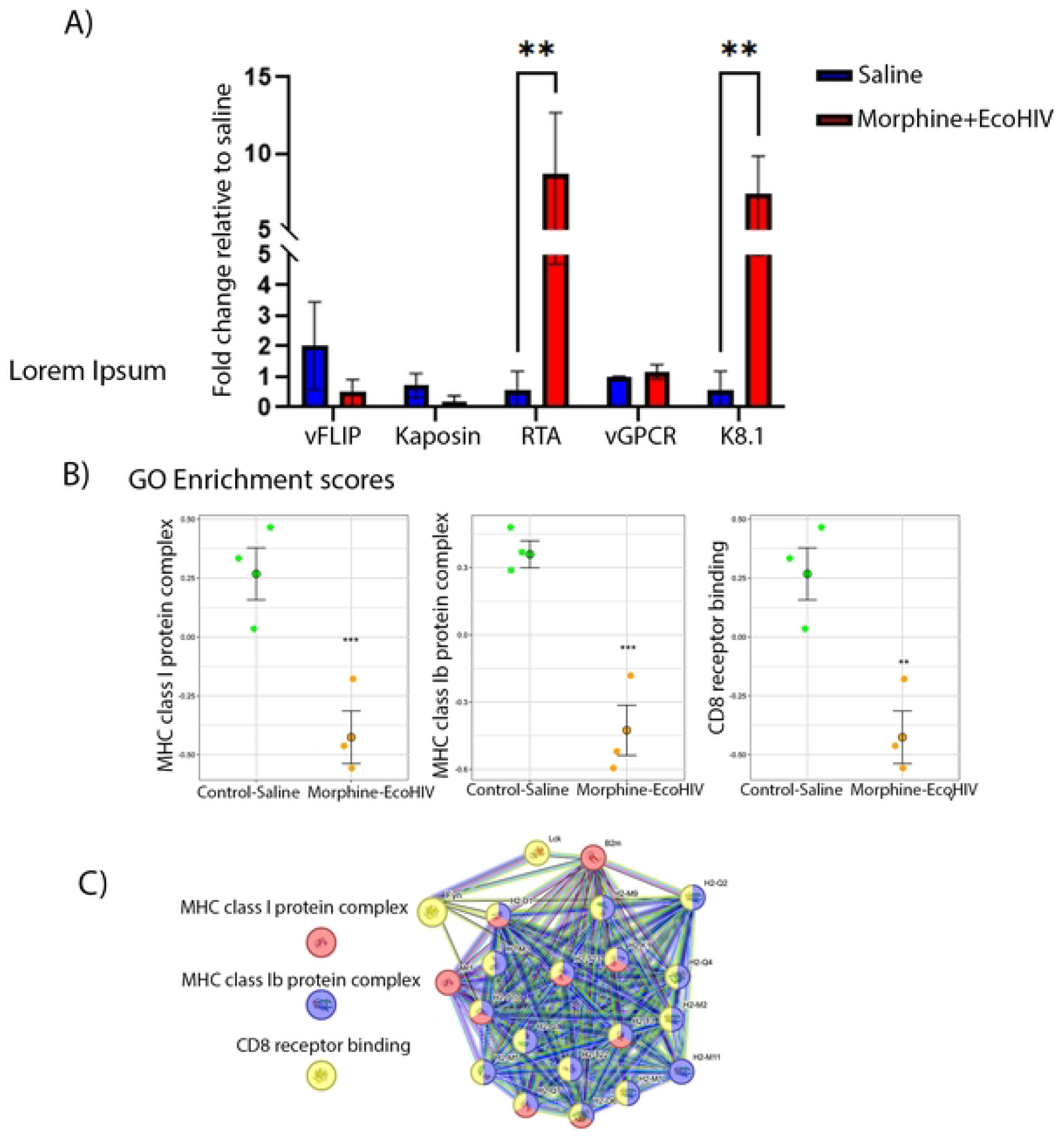
Mouse KS-like KSHV-positive tumors growing in Morphine+EcoHIV-treated immunodeficient mice engraft in immunocompetent mice. A) Schematic representation of the nude mice treatment and mECK36 cell injections. B) mECK36 KSHV-induced mouse KS-like tumor formation at seven weeks post-injection in untreated (blue) or Morphine+EcoHIV-treated (red) nude mice. C) Schematic representation of tumor transplantation. D) Engrafted tumors transplanted from untreated (blue) or Morphine+EcoHIV-treated (red) nude mice at seven weeks post-tissue implantation in untreated Balb/c mice.

To further evaluate the impact of opioid treatment and HIV infection on KSHV-infected tumor growth, and to determine if these treatments affected the tumor microenvironment, KS-like tumors from morphine+EcoHIV-treated or saline-treated (Untreated) nude mice were cut into small pieces (∼4mm^3^) and transplanted subcutaneously into untreated BALB/c immunocompetent mice (Figure 2C). These mice were monitored weekly for tumor formation. Tumor size (volume) was measured weekly until they reached approximately 2 cm^3^, at which point mice were euthanized.

In contrast to the results from mouse KS-like KSHV-positive tumors transplanted from saline-treated nude mice (Figure 2D, left panel), five of six mouse KS-like KSHV-positive tumors transplanted from morphine+EcoHIV-treated nude mice engrafted in untreated Balb/c immunocompetent mice (Figure 2D, right panel). This may indicate that the effect caused by the morphine+EcoHIV treatment could be partly due to tumor intrinsic changes, including changes in KSHV and host gene expression in the tumor and/or stromal cells.

### Mouse KS-like KSHV-positive tumors growing in Morphine+EcoHIV-treated immunodeficient mice showed increased KSHV lytic gene expression and downregulation of pathways involved in antigen presentation

Morphine+EcoHIV treatment increased KSHV lytic gene expression in mouse KS-like KSHV-positive tumors growing in nude mice, as shown by RT-qPCR analysis (Figure 3A). Moreover, we performed RNA-sequencing analysis on these tumors and compared them with those from saline-treated nude mice to better understand the mechanism behind the immunosuppressive effect. Our data showed that mouse KS-like KSHV-positive tumors from morphine+EcoHIV-treated nude mice exhibited downregulation of pathways related to MHC class I and Ib, as well as CD8 receptor binding, compared with tumors from saline-treated nude mice (Figure 3B). The most downregulated genes involved in these pathways included non-receptor tyrosine-protein kinases that play roles in T-cell differentiation (FYN and LCK), as well as genes associated with the major histocompatibility complex (MHC) class I and Ib (e.g., Mr1, B2m, H2-Q1, H2-M1)(Figure 3C)(Supplementary Table 1). Viruses have evolved several mechanisms to avoid recognition, many of which target the MHC-I antigen-processing pathway. KSHV encodes lytic proteins that remove MHC class I molecules from the cell surface, such as K3 and K5, so their increased expression upon treatment could play a role in this downregulation [28]. Moreover, HIV-1 renders the virus-infected cells less visible to CD8+ T cells through Nef-induced endocytosis of MHC-I from the cell surface[29]. These results point to possible mechanisms to explain why mouse KS-like KSHV-positive tumors growing in morphine+EcoHIV-treated nude mice engrafted into untreated Balb/c mice. A combination of KSHV lytic gene expression upregulation, downregulation of MHC class I, Ib antigen presentations, and CD8 receptor binding pathways, and the concomitant avoidance of KSHV-infected cell recognition by the immune system would explain the increased number of tumor engraftments in immunocompetent mice.

**Figure 3.**
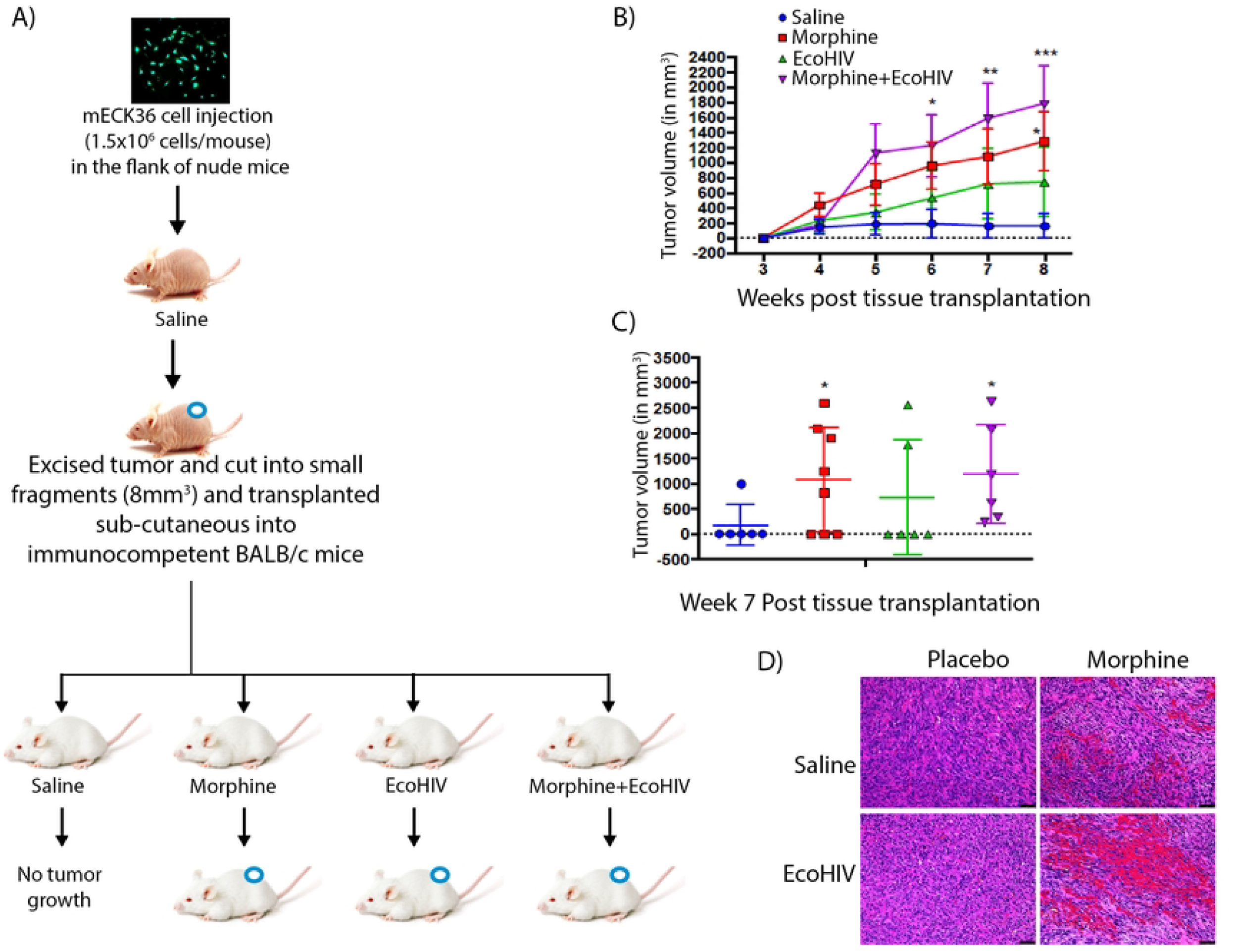
Mouse KS-like KSHV-positive tumors growing in Morphine+EcoHIV-treated immunodeficient mice showed increased KSHV lytic gene expression and downregulation of pathways involved in antigen presentation. A) Increase of KSHV lytic gene mRNA levels (by qRT-PCR) in KS-like tumors from morphine+EcoHIV-treated nude mice (red) compared to saline-treated nude mice (blue). (*p < 0.05; one-way ANOVA). B) RNA-sequencing analysis of KS-like KSHV-positive tumors growing in control-saline and morphine+EcoHIV-treated nude mice. KSHV-positive tumors from treated nude mice had down-regulation of pathways involved in MHC class I, Ib, and CD8 receptor binding when compared with tumors from saline-treated nude mice. (*p < 0.05; one-way ANOVA). C) Genes involved in the downregulated pathways.

### Morphine treatment and EcoHIV infection of immunocompetent mice allow mouse KS-like KSHV-positive tumor engraftment

To evaluate the effects of morphine and EcoHIV treatment on immunocompetent Balb/c mice, we designed a similar experiment in which we transplanted mouse KS-like KSHV-positive tumors that were growing in untreated nude mice. These tumors were cut into small pieces (∼4mm^3^) and transplanted subcutaneously into saline-treated, morphine-treated, and EcoHIV-infected BALB/c immunocompetent mice, either separately or in combination (Figure 4A). These mice were monitored weekly for tumor formation. Tumor size (volume) was measured weekly until they reached approximately 2 cm^3^, at which point mice were euthanized. (Figure 4B). As shown above (Figure 2D), only one of six mouse KS-like KSHV-positive transplanted tumors from nude mice successfully engrafted in saline-treated BALB/c mice. In contrast, morphine and EcoHIV treatment independently resulted in an increased number and size of engrafted tumors compared to saline-treated mice (Figure 4C). Remarkably, only BALB/c mice receiving morphine in combination with EcoHIV engrafted tumors in all transplanted mice, with an increased number and size compared to saline-treated mice and the treatments alone (Figure 4C). All tumors growing in immunocompetent Balb/c mice exhibited EGFP-positive cells, indicating that they are KSHVBac36-bearing tumors (Supplementary Figure 2). Moreover, H&E-stained sections of these tumors revealed the presence of inflammatory infiltrates and spindle cell/vasculature morphological alterations (Figure 4D).

**Figure 4.**
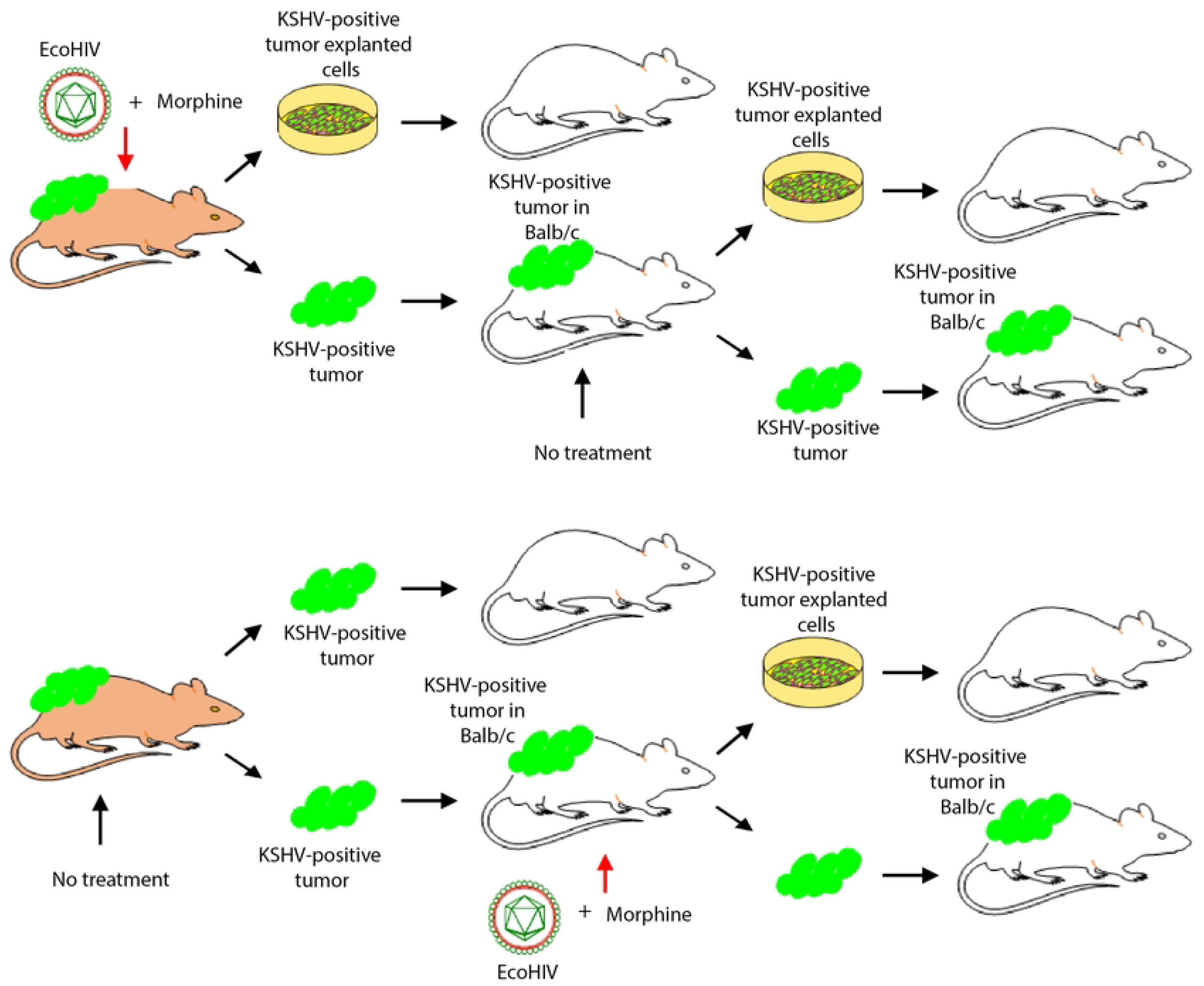
Morphine treatment and EcoHIV infection of immunocompetent mice allow mouse KS-like KSHV-positive tumor engraftment. A) Schematic representation of the KSHV-induced tumor formation in nude mice, subcutaneous tumor transplantation, and engraftment experiments in Balb/c mice. B) Subcutaneous tumor growth of transplanted mECK36 tumors in Balb/c mice treated with saline (blue), morphine (red), infected with EcoHIV (green), or the combination of morphine + EcoHIV (purple). Data indicate mean tumor size ± SEM (*p < 0.05; one-way ANOVA). C) Engrafted tumors at 7 weeks post tissue implantation in Balb/c mice. Data indicate mean tumor size ± SEM (*p < 0.05; one-way ANOVA). D) H&E-stained sections of KSHV-positive tumors growing in immunocompetent Balb/c mice.

When cells from mouse KS-like KSHV-positive tumors grown in Balb/c mice were explanted, disaggregated, and the tumor cells expanded, then re-injected into new Balb/c mice, they did not form tumors (Supplementary Figure 3A). However, when tumor fragments were directly implanted, they successfully formed tumors in untreated recipient Balb/c mice. This suggests that treatment-induced tumor characteristics are sustainable and can be propagated after passage from one mouse to another (Supplementary Figure 3B).

In conclusion, morphine and EcoHIV treatment enable and accelerate mouse KS-like KSHV-positive tumor engraftment in immunocompetent BALB/c mice. This effect is likely through the generation and recruitment of a stroma/cellular microenvironment with a dysregulated immune microenvironment permissive to tumor growth.

### Mouse KS-like KSHV-positive tumors growing in immunocompetent mice showed immune infiltration characteristic of KS lesions

Our newly designed model allows for the analysis of mouse KS-like KSHV-positive tumor immune environment/infiltration in immunocompetent mice. We performed immune infiltration analysis using RNA sequencing on KS-like KSHV-positive tumors growing in Balb/c mice. We defined the immune fraction profile using the Immune Cell Abundance Identifier for mouse (ImmuCellAI-mouse) algorithm[30], demonstrating significant enrichment of immune infiltrates in these KS-like KSHV-positive mouse tumors. We characterized the complexity of immune cells, including T cells, B cells, and monocytes (Figure 5A), all cellular populations described in human KS tissue biopsies[31]. We compared the immune infiltration of mouse KS-like KSHV-positive tumors growing in three different backgrounds: tumors from morphine+EcoHIV-treated donor nude mice but growing in control Balb/c mice (Saline); tumors from untreated donor nude mice but growing in morphine-treated Balb/c mice (Morphine); and tumors from untreated donor nude mice but growing in morphine+EcoHIV-treated Balb/c mice (Morphine+EcoHIV). We could not compare to tumors treated in vehicle only mice Balb/c, as these do not engraft. Importantly, mouse KS-like KSHV-positive tumors growing in EcoHIV-infected donor or recipient mice, in addition to the morphine, exhibited significantly lower immune infiltration scores compared to those in the Morphine-treated background with less B and T cells, as well as more macrophages, which are also abundant in KS tumors (Figure 5A and 5B). This downregulation in immune infiltration may explain the differences in tumor engraftment between these treatments, indicating that EcoHIV-infected backgrounds present an immune microenvironment that is more permissive to KSHV-positive tumor growth. ELISA determination of the anti-KSHV-antibody response was performed to assess reactivity to a panel of KSHV antigens, and we found that morphine alone treated Balb/c tumors displayed a robust anti-KSHV antibody response. In contrast, EcoHIV-treated tumors exhibited statistically less KSHV immune response (Figure 5C), correlated with the immune infiltration analysis.

**Figure 5.**
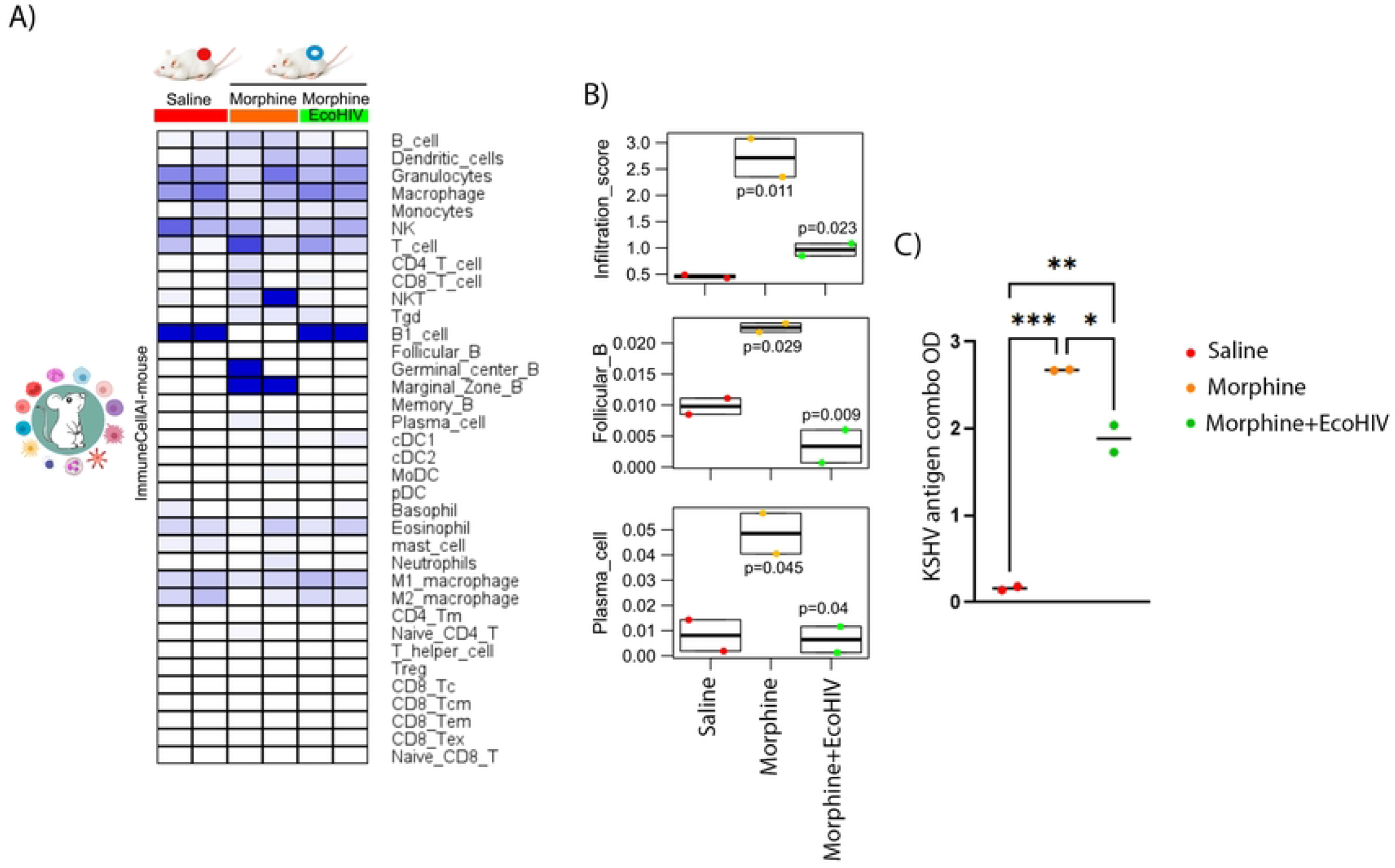
Mouse KS-like KSHV-positive tumors growing in immunocompetent mice showed immune infiltration characteristic of KS lesions. A) Heat map of enrichment scores of immune cell signature genes by RNA-Seq from KSHV-positive tumors growing in Control, morphine, and morphine+EcoHIV Balb/c mice. B) Boxplots show that infiltration decreases in KSHV-positive tumors growing in morphine+EcoHIV background. E) Anti-lytic and latent KSHV antigen combo ELISA in serum from Balb/c mice bearing KSHV-positive tumors. *P-values* were calculated by ANOVA.

## Discussion

KSHV-infected KS lesions primarily consist of latently infected cells and a few scattered lytically infected ones. Lytic genes can drive the development of KS lesions through paracrine mechanisms that promote proliferation and angiogenesis induced by viral genes such as the G protein-coupled receptor (vGPCR), K1, ORF45, or K15[1, 2, 32]. Two emerging therapeutic approaches related to these paracrine mechanisms in KS are mTORC1 inhibitors like Rapamycin[33, 34], which is highly effective only in transplant KS, suggesting immunomodulatory effects, and the use of Gleevec/Imatinib for AIDS-KS, aimed at targeting paracrine and autocrine activities mediated by c-kit and PDGFR, though it has shown limited efficacy in a subset of patients[35]. Current advances in novel therapies include target identification based on understanding KS’s pathogenesis and testing therapeutic modalities proven successful for other cancers, such as immunomodulators and cancer immunotherapy. These include Pomalidomide and PD-1/PD-L1 checkpoint inhibitors, which have shown partial success in a subset of individuals[8, 36]. A major gap in KS research is the fragmented evidence from in vitro infection, animal tumorigenesis, and clinical studies that yield numerous potential targets, which sometimes contradict one another. This underscores the need for an experimental system that unifies the pathogenesis of KS between experimental models and patients. This could cover the infection of KS oncogenic progenitors leading to tumorigenic outcomes that reflect clinical scenarios, helping to identify viral-host interactions driving KS tumors, which is critical for therapeutic interventions.

HIV/AIDS-KS results from multiple synergistic factors, which include KSHV-autonomous tumorigenesis, KSHV tumorigenesis-promoting elements such as inflammation and immunosuppressive mechanisms induced by HIV infection, including immune exhaustion, as well as immune senescence resulting from chronic HIV infection and inflammation. A prior study indicated that AIDS-KS is associated with markers of immunosenescence, characterized by an elevated frequency of CD4+ and CD8+ T-cells exhibiting an immunosenescence phenotype (CD57+ and CD28+) in KS cases compared to controls[37]. The current EcoHIV and morphine immunocompetent model of KSHV-permissive tumorigenesis can potentially recreate several of these factors. Although EcoHIV does not induce AIDS in murine models, our data suggest that it can, importantly, induce immunosuppression to anti-KSHV responses. Tumor engraftment was more effectively achieved with the combined treatment of morphine and EcoHIV. This is likely attributable to the fact that EcoHIV alone does not result in complete immunodeficiency; however, morphine is recognized to alter T cell function[13, 38]. Thus, both components are essential to replicate the proinflammatory and immunosuppressive consequences stemming from HIV infection in humans. Previous findings suggested that the combination of EcoHIV infection and morphine leads to an exacerbated inflammatory state relative to HIV infection alone, likely through influencing bacterial translocation and increasing inflammatory cytokines levels[25].

Our data indicate that the EcoHIV and morphine treatment enables mouse KS-like KSHV-positive tumors to grow in immunocompetent mice, likely through the generation and recruitment of an immunosuppressive stroma or cellular microenvironment. The growth of transplant tumors from morphine+EcoHIV-treated immunodeficient nude mice to untreated immunocompetent mice (Figure 6, upper panel) defines a system that could identify how these treatments induce an immunosuppressive microenvironment that helps overcome the rejection of KSHV-infected tumors and allows engraftment in immunocompetent mice. Our results point to viral lytic gene upregulation and the immune system’s avoidance of KSHV-infected cell recognition as possible mechanisms for overcoming rejection.

**Figure 6.**
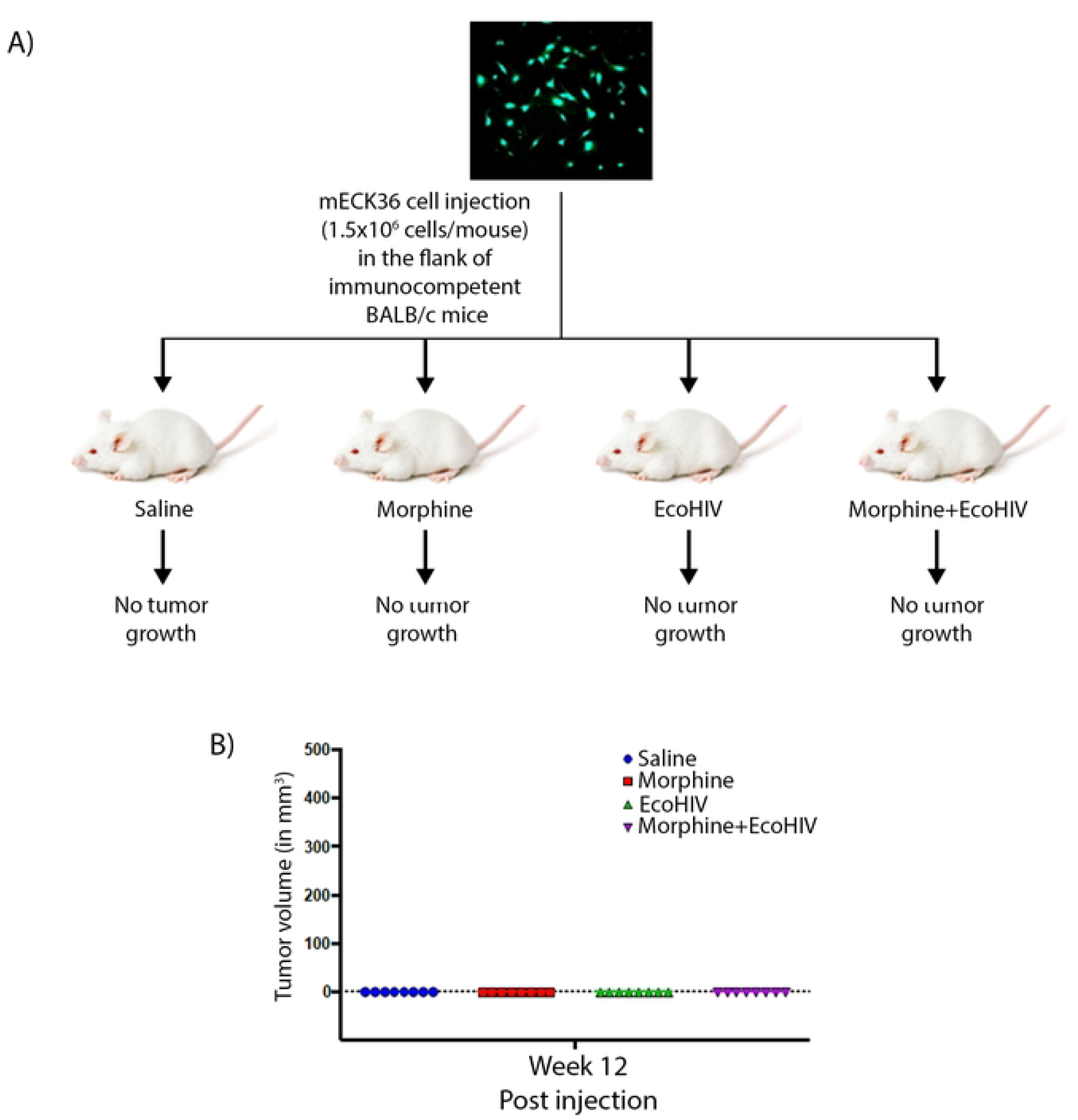
Two models of EcoHIV and morphine-induced immunosuppression to KSHV-infected tumor rejection in Balb/c mice: immunocompetent HIV/AIDS-KS tumor models. The upper panel defines a system that could allow the identification of how these treatments can help overcome the rejection of KSHV-infected tumors and allow engraftment to immunocompetent mice. The Lower panel is a system to study how these treatments induce immunosuppression in KSHV-infected tumors and allow their engraftment and growth in immunocompetent mice.

The model whereby mouse KS-like KSHV-positive tumors from untreated nude mice can be transplanted into immunocompetent mice after receiving EcoHIV and morphine treatments (Figure 6, lower panel) serves as a system to study how these treatments induce immunosuppression and allow engraftment and growth of KSHV-infected tumors in immunocompetent mice. More importantly, it constitutes an immunocompetent model of HIV/AIDS KS to study the role of HIV immunosuppression in facilitating the growth of KSHV-infected tumors. This model also represents a potentially powerful system to test immune-based anti-KS therapies and vaccines targeting KSHV, where an immune environment is required. These tumors growing in immunocompetent Balb/c mice will provide an excellent substrate to test immune-based anti-KS therapies, such as targeted antibodies or checkpoint inhibitors of clinical relevance for their potential use in AIDS-KS.

The possibility of having KSHV-driven tumors develop in immunocompetent mice due to EcoHIV infection presents a unique opportunity to create an AIDS-KS mouse model suitable for testing immune-based therapies and KSHV vaccines in a scenario that replicates important aspects of AIDS-KS. Both EcoHIV infection and morphine treatment in immunocompetent mice foster an immune environment that hinders anti-KSHV rejection and permits the growth of KSHV-infected tumors. This suggests that morphine likely acts through proinflammatory immune dysregulation mechanisms, while EcoHIV suppresses the immune response, probably due to a deficiency of CD4 T-cells, which are also crucial for helper-dependent CD8 T anti-gamma-herpesviral cytotoxicity as previously proposed[39]. We hypothesize that the development of these two models, as a set of HIV-positive and HIV-negative KSHV tumor systems in Balb/c, forms a novel and unique HIV/AIDS KSHV tumorigenesis system in Balb/c immunocompetent mice to evaluate established immune-based therapies against KS-like tumors, including vaccines. More importantly, this represents an immunocompetent yet immunodysfunctional mouse model of HIV/AIDS associated KS to investigate the role of HIV in facilitating the growth of KSHV-infected tumors.

The demonstration that treatment with EcoHIV and morphine allows the establishment of KSHV-infected tumors, which can be serially passaged in immunocompetent mice, is the first demonstration that a preclinical model can be developed representing KS-like tumors in the context of an immune response, albeit impaired, as is the case in people with HIV and KS. These first-in-kind murine models incorporating a biologically relevant mouse HIV infection tool make our models unique and original, with the potential to provide novel mechanistic and pathobiological insights and drive the development of novel therapies that, for the first time, will allow preclinical testing of immune-based therapies and vaccines for KS.

### Materials and Methods Animal

For all our studies in immunocompetent mice, we used 8-12-week-old male BALB/c mice. The use of BALB/c is appropriate for our studies because our mouse KS model is based on the BALB/c backbone[20], and they are also susceptible to EcoHIV infection[25]. All mice were kept under standard conditions and used under protocols approved by the Institutional Animal Care and Use Committee of the University of Miami.

### Morphine treatment

Mice were implanted with either a placebo or a slow-release morphine pellet 24 h before infection with EcoHIV. Pellets were generously provided by NIH/NIDA. The mice were implanted with either 25 mg or 75 mg morphine pellets using a protocol that would increase the release of morphine every 7 days to avoid tolerance. Escalating doses in the order will be used: 1× 25 mg, 1× 75 mg, 2× 25 mg, and 2× 75 mg based on a study that showed 2× 25 mg morphine pellets resulted in higher morphine release than 1× 75 mg pellet, likely due to increased surface area with having two pellets[40]. The first dose was given (implant at day 0, exposed 0–7 days), 25mg morphine pellet as in short-term experiments. In the second week (implant at day 7, exposed 7– 14 days), mice were implanted with a 75mg morphine pellet. In the third week (implant at day 14, exposed 14–21 days), mice were pelleted with 2×25 mg pellets, and in the final week (implant at day 21, exposed 21–28 days) with 2×75 mg pellets.

### EcoHIV Infection

The EcoHIV, a modified HIV that can infect mouse cells, was used in this study. EcoHIV provides a useful avenue for studying HIV and drugs of abuse, where other models have fallen short. This model provides flexibility for looking at different opioid paradigms concerning HIV infection, in addition to having the ability to utilize different genetic backgrounds, such as MORKO mice, which are not available when using humanized mice. The construct EcoHIV/NL4-3-GFP was used in this study, which was constructed on the backbone of HIV-1/NL4-3 and expresses enhanced green fluorescence protein (EGFP) as a marker. The virus is called EcoHIV-GFP. The virus was propagated in HEK293TN cells as previously described[25], and the virus was estimated by p24 ELISA (ZeptoMetrix Corporation). A total of 0.5-1 mL of either saline or 1×106 pg. p24 equivalent EcoHIV was injected into the intraperitoneal cavity (i.p.).

### KS mouse model

It has been shown that mECK36 is a biologically sensitive animal model of virally induced KS. Transfection of KSHV Bacterial Artificial Chromosome (KSHVBac36) into mouse bone marrow endothelial-lineage cells generates a cell (mECK36) that forms KS-like KSHV-positive tumors in immunodeficient mice[20].

### KSHV tumorigenesis

Tumors were induced by subcutaneous (s.c) injection of 1×10^6^ mECK36 cells to untreated or morphine-treated + Eco-HIV-infected nude mice (n=6). mECK36 formed solid tumors 3 weeks after injection. To assess the effects of morphine treatment and EcoHIV infection on tumorigenesis, we determine the growth of the tumor by daily measurements with a caliper and applying the formula volume = length x width2 × 0.52. mECK36 tumors were allowed to grow until they reached a volume of ∼2 cm[20]. All mice were sacrificed, and complete necropsies were performed. The blood was collected, and various tissues/organs were isolated aseptically for different analyses and stored at -80 °C.

### Immunocompetent mice tumor engraftment

mECK36 KS-like KSHV-positive tumors in nude mice were cut into small pieces (∼4mm^3^) and transplanted subcutaneously in the flank of untreated/control, morphine-treated, EcoHIV-infected BALB/c immunocompetent mice separately or in combination. Mice were observed at weekly intervals for tumor formation. Tumor size (volume) was measured weekly until mice were euthanized. Tumors were allowed to grow until they reached a volume of ∼2 cm[20]. All mice were sacrificed, and complete necropsies were performed. The blood was collected, and various tissues/organs were isolated aseptically for different analyses and stored at -80 °C.

### RNA sequencing

RNA was isolated and purified from KS-like KSHV-positive tumors using the RNeasy Mini Kit (Qiagen), following the manufacturers protocol for optimal RNA extraction. The quality and integrity of RNA were assessed using the Agilent Bioanalyzer 2100, and concentrations were quantified with a NanoDrop spectrophotometer to ensure suitability for downstream sequencing. A total of three biological replicates per condition were included to ensure statistical rigor and biological reproducibility. The purified RNA samples were sent to Novagen for library preparation and high-throughput RNA sequencing on the Illumina platform. Sequencing reads were aligned to the mouse reference genome using STAR aligner, and gene-level counts were generated for downstream analysis.

Differential gene expression analysis was conducted using the DESeq2 package in R/Bioconductor. Raw counts were normalized using DESeq2’s internal method, which adjusts for differences in sequencing depth and RNA composition across samples. Statistical significance was assessed using the Wald test, and p-values were adjusted for multiple comparisons using the Benjamini-Hochberg procedure to control the false discovery rate (FDR). Genes with an adjusted p-value (FDR) < 0.05 were considered significantly differentially expressed.

For pathway-level interpretation, we employed Gene Set Variation Analysis (GSVA) to evaluate the enrichment of predefined gene sets in an unsupervised manner across individual samples. This approach allowed for the identification of pathway activity differences between experimental groups. Additionally, the STRING database was used to analyze protein-protein interaction networks, providing further insights into functional relationships among differentially expressed genes.

The Immune Cell Abundance Identifier for mouse resource (ImmuCellAI-mouse, https://guolab.wchscu.cn/ImmuCellAI-mouse/) was employed to estimate the abundance of 36 immune cells based on RNA sequencing gene expression profile obtained from each sample.

### Real-Time Quantitative PCR (RT-qPCR)

RNA was isolated and purified from KS-like KSHV-positive tumors as mentioned above. To remove DNA, samples were treated with RNAse-free DNase I (QIAGEN) on columns for 25 minutes at room temperature. RNA was reverse-transcribed into cDNA using ImProm-II Reverse Transcriptase (Promega) according to the manufacturer’s protocol. Viral and host messenger RNA (mRNAs) were amplified using specific primers diluted in the SYBR green PCR master mix (Quanta Biosciences). For detection, we used the Atila machine and software. Nonreverse transcriptase and water controls were used to confirm the samples’ absence of viral DNA and contamination. Actin was used as a housekeeping gene to perform the ΔΔCT method. The expression of viral genes was normalized to the actin CT value, and the difference between viral CT values with actin was considered the ΔCT value. The obtained ΔCT values were normalized to a given control sample (ΔΔCT value). The fold target gene expression is given by the formula: 2–ΔΔCT.

### KSHV ELISA

Anti-KSHV antibodies were determined on all samples using an ELISA composed of KSHV antigen-coated plates. Diluted test sera and control sera (100 µl at 1:200 in assay buffer) were pipetted into the wells and incubated at 37°C for 60 minutes. Positive and negative controls were run in duplicate on each plate. After incubation, the wells were emptied, and the plate was washed five times with 1x wash buffer. Residual liquid was removed by sharply tapping the plate on absorbent paper. 100 uL of diluted Anti-Mouse IgG Enzyme Conjugate was added to each well, mixed for 30 seconds, and incubated at 37 °C for 1 hour. The wash procedure was repeated as described above. 100 uL of TMB (3,3’,5,5’-tetramethylbenzidine) substrate was dispensed into each well and mixed gently. The reaction was allowed to develop at room temperature (18-22 °C) for 20 minutes. 100 uL of stop solution was added to each well to halt the enzymatic reaction. Absorbance was measured at 450 nm using a microplate reader. The optical density (OD) was used to determine the presence or absence of anti-HHV-8 antibodies in the samples.

## Acknowledgments

Dr. Enrique A. Mesri, an inspiring scientist, a great mentor, and a friend, sadly passed away before sending the manuscript. For his dedication and contributions to the KSHV and viral oncology field, we will always be in debt to him.

## Figure legends

**Supplementary Figure 1**. A) Schematic representation of untreated Balb/c mice injections with KSHV-positive mECK36 cells. B) Tumor growth in Balb/c mice.

**Supplementary Figure 2**. GFP-positive isolated tumor cells from Mouse KS-like KSHV-positive tumors growing in Balb/c mice.

**Supplementary Figure 3**. A) Schematic representation of the animal injections with isolated KSHV-positive tumor cells from Balb/c mice. B) Schematic representation of the animal transplants with Mouse KS-like KSHV-positive tumor fragments from Balb/c mice.

